# Complete genome sequence of a divergent strain of Tibetan frog hepatitis B virus associated to concave-eared torrent frog *Odorrana tormota*

**DOI:** 10.1101/506550

**Authors:** Humberto J. Debat, Terry Fei Fan Ng

**Affiliations:** Instituto de Patología Vegetal, Centro de Investigaciones Agropecuarias, Instituto Nacional de Tecnología Agropecuaria (IPAVE-CIAP-INTA), X5020ICA, Córdoba, Argentina.; College of Veterinary Medicine, University of Georgia, Athens, Georgia, USA 30602.

**Keywords:** dsDNA virus, Hepadnavirus, amphibian virology, frog virus, Odorrana tormota, virus discovery

## Abstract

The family *Hepadnaviridae* is characterized by partially dsDNA circular viruses of approximately 3.2 kb, which are reverse transcribed from RNA intermediates. Hepadnaviruses (HBVs) have a broad host range which includes humans (Hepatitis B virus), other mammals (genus *Orthohepadnavirus*), and birds (*Avihepadnavirus*). HBVs host specificity has been expanded by reports of new viruses infecting fish, amphibians, and reptiles. The tibetan frog hepatitis B virus (TFHBV) was recently discovered in *Nanorana parkeri* (Family *Dicroglossidae*) from Tibet. To increase understanding of hepadnavirus in amphibian host, we identified the full-length genome of a divergent strain TFHBV-Ot associated to the concave-eared torrent frog *Odorrana tormota* (Family *Ranidae*) from China by searching deep sequencing data. TFHBV-Ot shared the genomic organization and a 76.6% overall genome nucleotide identity to the prototype TFHBV associated to *N. parkeri* (TFHBV-Np). TFHBV-Ot amino acid pairwise identity with TFHBV-Np predicted gene products ranged between 63.9% and 77.9%. Multiple tissue/organ specific RNAseq datasets suggest a broad tropism of TFHBV including muscles, gonads and brains. In addition, we provide for the first time evidence of virus derived small RNA from an amphibian hepadnavirus, tentatively enriched in 19-20 nt species and cytidine as first base. The results presented here expand the genetic diversity and the host range of TFHBV to *Ranidae* frogs, and warrant investigation on hepadnaviral infection of amphibian brains.

## Main

The family *Hepadnaviridae* is characterized by viruses with partially dsDNA circular genomes of ca. 3.2 kb, which are reverse transcribed from RNA intermediates. Hepadnaviruses (HBVs) have a broad host range which includes humans (Hepatitis B virus), other mammals (genus *Orthohepadnavirus*) and birds (*Avihepadnavirus*). More recently, HBVs have been reported to infect fish, amphibians, and reptiles. The Tibetan frog hepatitis B virus (TFHBV) was discovered recently in *Nanorana parke*i (Family *Dicroglossidae*) from Tibet, expanding the diversity of hepadnaviruses to amphibians: a neglected virus host [1]. The emergent clade of amphibians and reptile HBVs has been tentatively clustered as “herpetohepadnaviruses” within the *Hepadnaviridae* family [2]. The concave-eared torrent frog (*Odorrana tormota*) distribution is restricted to the norther region of the costal Zhejiang province of China. Concave-eared male frog have a large and distinctive call repertoire, which has been linked to ultrasonic communication [3]. While exploring by BLASTX searches a transcriptome dataset of *O. tormota,* oriented to unravel molecular mechanisms of concave-eared frog growth and development [4] (BioProject PRJNA437724), we found a 7.9 kb long contig of a hepadnavirus-like sequence (E-value = 0). This contig was flanked by identical regions at either end, typical of an assembled sequence from a circle genome. Using dotpot as implemented in http://www.bioinformatics.nl/emboss-explorer/, the identical region was removed to construct a circle genome of 3,144 base pairs. In addition, the 1,412,314,266 raw 150 bp reads from PRJNA437724 were used to polish the contig using the Geneious v8.1.9 platform (Biommaters, USA) map to reference tool with low sensitivity parameters. The genome was supported by 40,719 reads (mean coverage = 2,348X, minimum cov. 190X, maximum cov. 6,959X) (**Table 1, Supp. Figure 1.A**). The predicted circular nature of the virus sequence was further confirmed by identifying overlapping read at both sequence termini, supported by >5,000 reads simultaneously covering both terminal regions (**Supp. Figure 1.B**). In addition, 14 small RNA datasets of the same bioproject [4], derived from diverse tissue/organs of six concave-eared torrent frogs were assessed. Interestingly, in five of the libraries of two animals we detected virus-derived small RNAs, encompassing over 5,804 reads (**Supp. Table 1**), ranging from 18-37 nt long. The specific landscape derived from sRNA mapping to the virus sequence suggested an asymmetrical distribution or reads and the presence of tentative hot-spots which could be associated to eventual regions of the virus RNA more prone to generate vsRNAs (**Supp. Figure 1.C**). Additional analyses of the small RNA datasets were conducted, and we observed that overall frog small RNAs are enriched in 21-22 nt species, as observed by Shu et al specifically for microRNA species [4], and starting mostly with a Uridine as first base (**Supp. Figure 2 A-B**). On the other hand, virus derived sRNAs appear to be enriched in 19-20 nt species, with a more diverse set of first bases, marginally enriched in Cytidine (**Supp. Figure 2 A-B**). Given the relatively low number of virus derived sRNA species detected, further experiments should assess if the tendencies observed here are supported, or derived from low sample size. To our knowledge, this is the first time that hepadnavirus derived small RNAs are described in any amphibian, suggesting that the virus RNA is recognized by the RNA interference machinery of the frog host, which might be related to the induction of a molecular response to virus infection.

**Table 1.** Tibetan frog hepatitis B virus derived reads in diverse NGS RNAseq datasets obtained from total RNA of dissected tissue/organs from *Odorrana tormota*. Data is paired 150bp reads obtained with HiSeq X Illumina instrument. Viral reads less than 100 would be treated as typical sequencing bleed over.

| Sample id | Run | BioSample | Experiment | Sex | Tissue | Total reads | Virus reads | Virus RPM |
| --- | --- | --- | --- | --- | --- | --- | --- | --- |
| OT1FB | SRR6896130 | SAMN08683453 | SRX3845984 | female | Brain | 50925040 | 1846 | 36.2 |
| OT1FE | SRR6896129 | SAMN08683456 | SRX3845985 | female | Auricularis muscle | 50767620 | 10317 | 203.2 |
| OT1FM | SRR6896120 | SAMN08683463 | SRX3845994 | female | Hindlimb muscle | 50832762 | 2374 | 46.7 |
| OT1FS | SRR6896132 | SAMN08683459 | SRX3845982 | female | Ovaries | 50809536 | 173 | 3.4 |
| OT1MB | SRR6896119 | SAMN08683466 | SRX3845995 | male | Brain | 51520470 | 40 | 0.8 |
| OT1ME | SRR6896122 | SAMN08683469 | SRX3845992 | male | Auricularis muscle | 51145732 | 33 | 0.6 |
| OT1MM | SRR6896144 | SAMN08683477 | SRX3845970 | male | Hindlimb muscle | 50581552 | 30 | 0.6 |
| OT1MS | SRR6896127 | SAMN08683472 | SRX3845987 | male | Testis | 51398962 | 43 | 0.8 |
| OT2FB | SRR6896131 | SAMN08683454 | SRX3845983 | female | Brain | 51227736 | 2459 | 48.0 |
| OT2FE | SRR6896134 | SAMN08683457 | SRX3845980 | female | Auricularis muscle | 51446264 | 17031 | 331.0 |
| OT2FM | SRR6896121 | SAMN08683464 | SRX3845993 | female | Hindlimb muscle | 50894374 | 4673 | 91.8 |
| OT2FS | SRR6896133 | SAMN08683460 | SRX3845981 | female | Ovaries | 50794092 | 398 | 7.8 |
| OT2MB | SRR6896124 | SAMN08683467 | SRX3845990 | male | Brain | 50484980 | 34 | 0.7 |
| OT2ME | SRR6896123 | SAMN08683470 | SRX3845991 | male | Auricularis muscle | 51127928 | 38 | 0.7 |
| OT2MM | SRR6896143 | SAMN08683478 | SRX3845971 | male | Hindlimb muscle | 50731628 | 0 | 0.0 |
| OT2MS | SRR6896141 | SAMN08683473 | SRX3845973 | male | Testis | 50758148 | 4 | 0.1 |
| OT3MF | SRR6896138 | SAMN08683476 | SRX3845976 | male | Skin | 54287526 | 0 | 0.0 |
| OT3MS | SRR6896139 | SAMN08683475 | SRX3845975 | male | Testis | 51007646 | 0 | 0.0 |
| OT4FB | SRR6896128 | SAMN08683455 | SRX3845986 | female | Brain | 54737374 | 0 | 0.0 |
| OT4FE | SRR6896135 | SAMN08683458 | SRX3845979 | female | Auricularis muscle | 55143698 | 3 | 0.1 |
| OT4FF | SRR6896137 | SAMN08683462 | SRX3845977 | female | Skin | 50844636 | 0 | 0.0 |
| OT4FM | SRR6896118 | SAMN08683465 | SRX3845996 | female | Hindlimb muscle | 54540910 | 0 | 0.0 |
| OT4FS | SRR6896136 | SAMN08683461 | SRX3845978 | female | Ovaries | 55815918 | 4 | 0.1 |
| OT4MB | SRR6896125 | SAMN08683468 | SRX3845989 | male | Brain | 55649520 | 153 | 2.7 |
| OT4ME | SRR6896126 | SAMN08683471 | SRX3845988 | male | Auricularis muscle | 54601460 | 645 | 11.8 |
| OT4MM | SRR6896142 | SAMN08683479 | SRX3845972 | male | Hindlimb muscle | 54529624 | 214 | 3.9 |
| OT4MS | SRR6896140 | SAMN08683474 | SRX3845974 | male | Testis | 55709130 | 207 | 3.7 |

This hepadanvirus in concave-eared torrent frog is a novel TFHBV strain Ot (TFHBV-Ot), as it shares a 76.6% overall genome nucleotide identity to the prototype TFHBV-Np. We analyzed the virus sequence for ORFs and conserved amino acid domains, using established tools (ORFfinder, https://www.ncbi.nlm.nih.gov/orffinder/; conserved domain database search tool, https://www.ncbi.nlm.nih.gov/Structure/cdd/wrpsb.cgi and compared with the prototype TFHBV reference. The 3,144-bp TFHBV-Ot sequence contains a typical hepadnaviral organization (**Figure 1.A**), consisting in three major overlapping ORFs associated with the Core (PreC/C), Polymerase (P) and Surface (PreS/S) genes, encoding for capsid subunits, viral DNA polymerase, and surface protein, respectively. As in the prototype TFHBV, the virus sequence presented no evidence of an X protein, which has been associated with mammal-infecting orthohepadnaviruses [6]. Multiple sequence alignments (MSA) using MAFTT v7.017 [5] (E-INS-i algorithm, BLOSUM 62 scoring matrix) showed that the predicted gene products presented a 77.9% (C), 72.1% (P), and 63.0% (S) aa identity to that of the prototype TFHBV associated to *N parkeri* (TFHBV-Np) (**Figure 1.B**).

**Figure 1.**
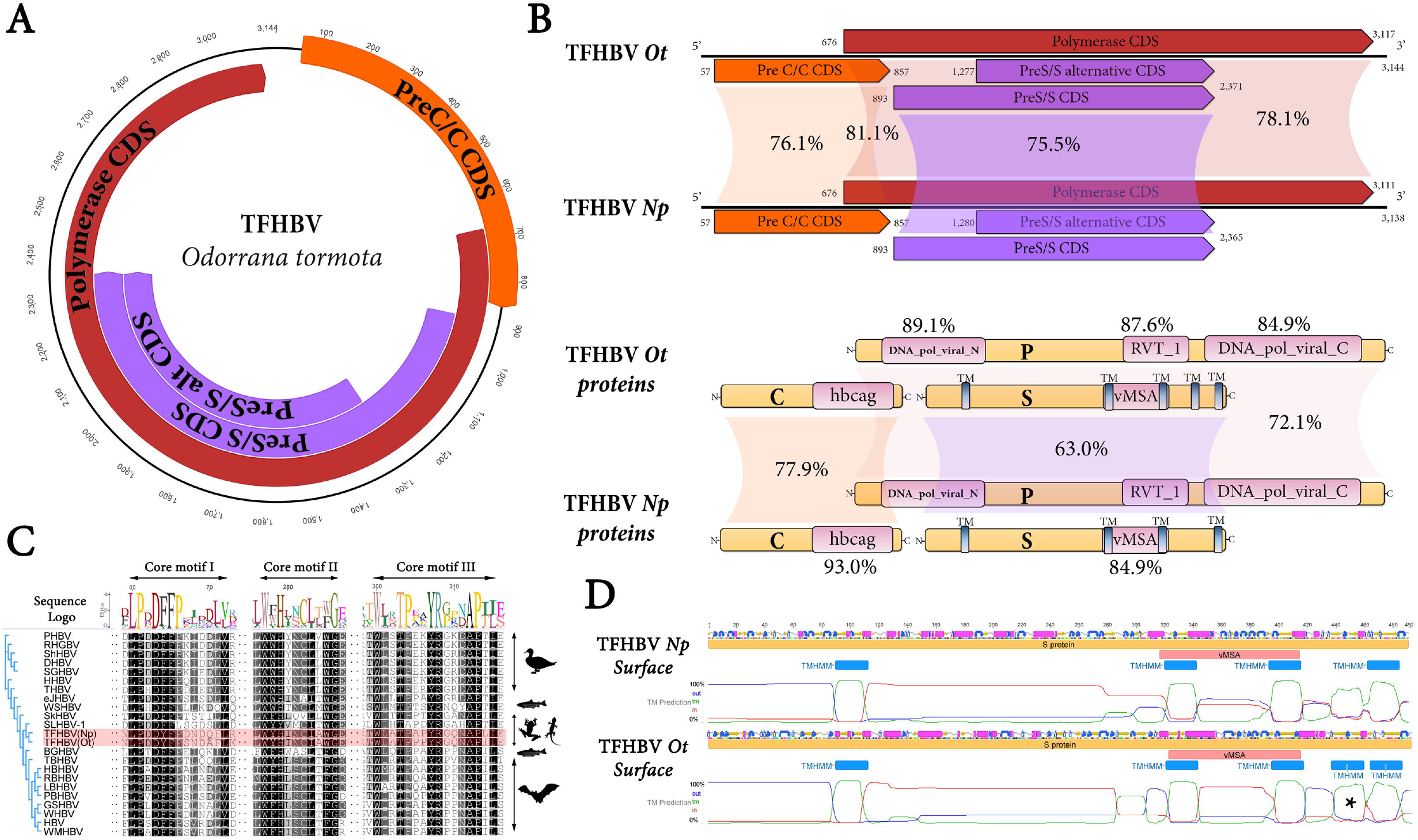
Molecular characterization of tibetan frog hepatitis B virus associated to *Odorrana tormota* (TFHBV-Ot) (**A**) Genome graph depicting predicted gene products of TFHBV-Ot. The predicted coding sequences are shown in orange (PreC/C), red (P) and purple (S) arrowed rectangles. (**B**) Gene products architecture and comparison of TFHBV-Ot and Nanorana parkeri associated TFHBV. Start and end coordinates of each gene product are indicated, and nt pairwise identity are shown as percentage values (upper panel). Curved yellow rectangles represent each predicted proteins and conserved domains are shown in pink. aa pairwise identity is represented in percentage values for the overall predicted proteins, or for each specific domain (lower panel). Abbreviations: DNA_pol_viral_N/C, N/C-terminal domain of the viral DNA polymerase; RVT_1, Reverse transcriptase; vMSA, Major surface antigen from hepadnavirus; hbcag, Hepatitis core antigen. (**C**) Multiple aa alignment of C proteins showing the conserved core motifs I-II-III. Silhouettes illustrate representative host organism of the respective virus. TFHBV are indicated with red highlighting. For virus abbreviations please refer to **Supp Table 2.** (**D**) Secondary structure of TFHBV surface protein as predicted with Emboss garnier (coils in grey, alpha helix pink, turns in blue arrows and beta strands in yellow arrows). Transmembrane helices in proteins predicted by TMHMM Server v. 2.0 are indicated with blue rectangles. Asterisk shows an additional transmembrane site at the C-terminal region of TFHBV-Ot S.

Our comparative analysis between TFHBV-Ot and the prototype genome revealed common features for this virus species. The first CDS (PreC/C) extends between the 57-857 nt coordinates, encoding a 30.6 kDa 266 aa phosphoprotein involved in assembly of subviral capsids [1]. TFHBV-Ot shares with TFHBV-Np (also 266 aa and 31.3 kDa) the specific domains of the C protein at equilocal positions, including the conserved core motifs I-II-III [1] (**Figure 1.C**). TFHBV-Ot contains a hepatitis core antigen domain (hbcag, pfam00906, E-value = 3.74e-22), with 93% AI (amino acid identity) to the prototype strain at the capsid core domain region (166-208 aa coordinates, InterPro id IPR036459). Both TFHBV strains shared the arginine rich core C-terminal region which has been associated to nuclear transport signal for pre-genome encapsidation [8]. The PreC/C CDS overlaps by 182 nt with the P CDS which is located between the 676-3,117 nt coordinates. This gene encodes a 92.3 kDa viral DNA polymerase. Similar to the prototype, TFHBV-Ot lacks the orthohepadnavirus-specific expansion in both the N-terminal of DNA polymerase domain and reverse transcriptase domain (**Supp. Figure 2**). While overall AI between TFHBV-Ot and TFHBV-Np P was 72.1%, AI reaches as high as 89.1% at functional domains such as the N-terminal of the polymerase (**Figure 1.B**). Prototype TFHBV-Np was found to potentially encode two alternative PreS/S start codon positions [1], the same configuration was detected in TFHBV-Ot as well. Even though TFHBV-Ot appears to share similar S folding and topology to the prototype, we identified an additional transmembrane signal at the C region of the longer S, absent in TFHBV-Np, which could potentially form an extra loop in the ER lumen (**Figure 1.D**; TMHMM v. 2.0 tool, http://www.cbs.dtu.dk/services/TMHMM/).

In general, hepadnaviruses are characterized by narrow host specificity and marked hepatotropism. Nevertheless, some avihepadnaviruses have been detected in extra-liver organs, such as pancreas, kidney and spleen [9]. There are no reports assessing the tropism of amphibian viruses in general, or hepadnaviruses in particular. Here, we analyzed multiple RNAseq datasets of *O. tormota* in order to detect potential virus derived RNAs, which could be considered as indirect evidence of a tentative tropism of TFHBV. The bioproject PRJNA437724 is composed of high throughput sequencing in HiSeq X Ten platform (Illumina, USA) of RNA extractions of dissected tissues/organs of four adult male and three female *O. tormota* frogs from the Anhui Province of China [4]. The raw 150 bp reads from each library of this bioproject were mapped to TFHBV-Ot genome using the Geneious v8.1.9 map to reference tool with low sensitivity parameters. Interestingly, at least three of the seven individual frogs presented strong RNA evidence of TFHBV (**Table 1**). In descending order of viral reads, the virus were detected consistently in auricularis and hindlimb muscles, brain, and ovaries. In addition the frogs presenting higher virus RNA titters (Females OT1 and OT2), when assessed by sRNA sequencing, were also found to present virus derived sRNAs from both hindlimb muscles and brain libraries (**Supp. Table 1**). While further direct detection experiments are needed to support these findings, TFHBV might have a broad tropism in *O. tormota* frogs. In healthy animals, the brain is a pristine organ protected from infection by the blood-brain barrier. The presence of detectable viral RNA in the brain tissue of the three positive frogs suggest potential central nervous system infection. Even though sample contamination from blood to brain during collection cannot be ruled out, this finding warrants further investigation on potential hepadnaviral pathogenicity in brain tissue of amphibians.

To investigate the evolution of TFHBV, phylogenetic analysis were performed using MAFFT alignments (BLOSUM 62 scoring matrix; E-INS-i, L-INS-i, and G-INS-i algorithm, respectively) and maximum likelihood FastTree v2.1.10 http://www.microbesonline.org/fasttree/trees (JTT aa evolution model, CAT approximation, local support values with Shimodaira-Hasegawa test) of P, C, and S proteins of TFHBV-Ot and reported hepadnaviruses (**Figure 2.A-C**). The resulting trees unequivocally clustered both TFHBV strains into a monophyletic clade of frog and reptiles viruses (dubbed herpetohepadnaviruses [2]), including skink hepatitis B virus and spiny lizard hepatitis B virus, which were discovered recently in public sequence databases of *Saproscincus basiliscus* and *Sceloporus adleri*, respectively [2]. Hepadnaviruses were suggested to harbor the most frequent virus-host co-divergence levels of any animal virus family [7]. The relatedness of the two TFHBV strains with considerably close host geographic proximity (*Nanorana parkeri* frog from Tibet and *Odorrana tormota* frog from Anhui Province of China) suggested that the evolution could be the result of virus–host co-divergence. However, in depth analysis is precluded by the limited number of amphibian hepadnavirus known. Future studies should unveil whether TFHBV might be linked to additional frog species and whether any pathogenic effect, such as brain disease, is associated to hepadnavirus infection on amphibian hosts.

**Figure 2.**
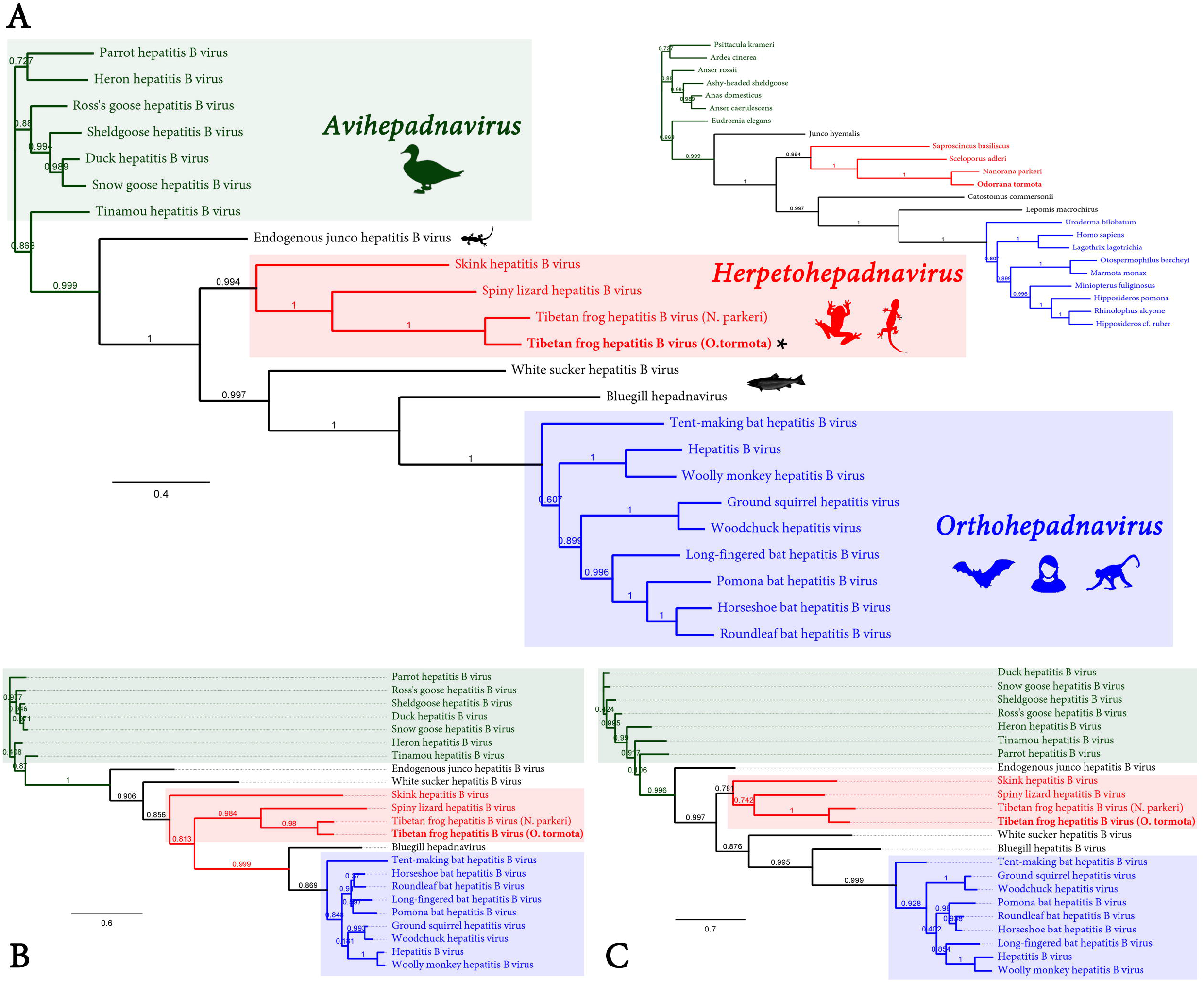
Phylogenetic insights of TFHBV-Ot based on MAFFT alignments and maximum likelihood trees generated with P (**A**), C (**B**), and S proteins (**C**) of TFHBV-Ot and reported hepadnaviruses. The scale bar indicates the number of substitutions per site. Node labels indicate FastTree support values. Silhouettes illustrate representative host organism of the respective virus. Viruses clustering into genera *Avihepadnavirus, Orthohepadnavirus* and the putative herpetohepadnavirus clade are indicated with green, blue and red rectangles, respectively. The right inset of panel A mirrors the phylogenetic tree on the left indicating the reported host of each virus.

## Data availability

The genome sequence reported here has been deposited in GenBank under the accession number MH700450.

## Supporting information

TFHBV-Ot virus sequence

## Acknowledgements

HJD would like to thank Benjamin Neuman for helpful discussions and insightful comments.

## Ethical approval

This article does not contain any studies with human participants or animals performed by any of the authors.

---

**Supplementary Figure 1.**
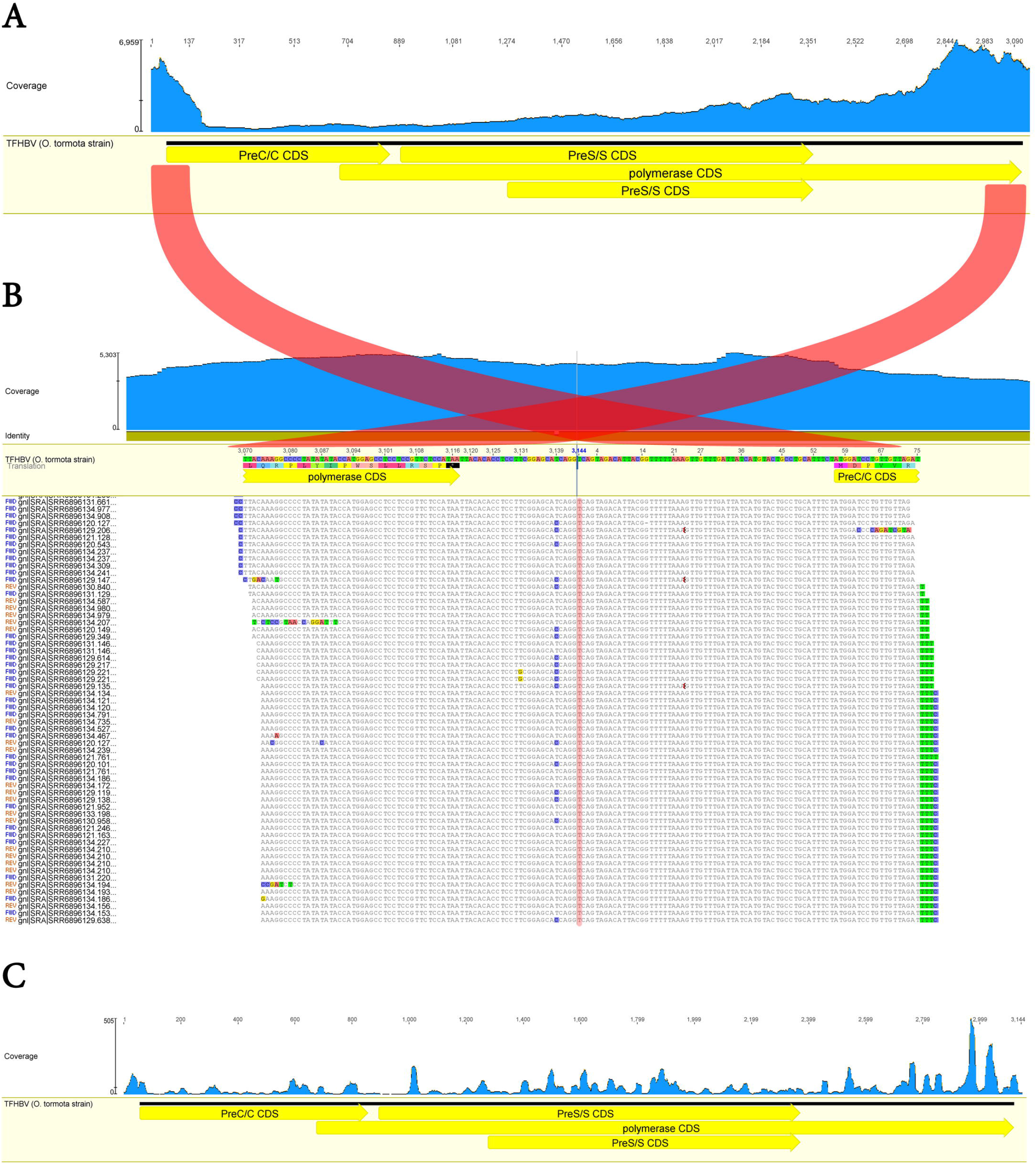
Coverage landscape per position obtained by mapping of total RNA virus derived reads to TFHBV-Ot (**A**) or small RNA virus derived reads (**C**). Scale represents nt coverage per position. In (**B**) total RNA reads were mapped to the terminal region of TFHBV-Ot and flanking 150 bp showing high coverage of overlapping reads mapping to both terminal regions, supporting the circular nature of the virus sequence. Position one of TFHBV-Ot is highlighted in red.

**Supplementary Figure 2.**
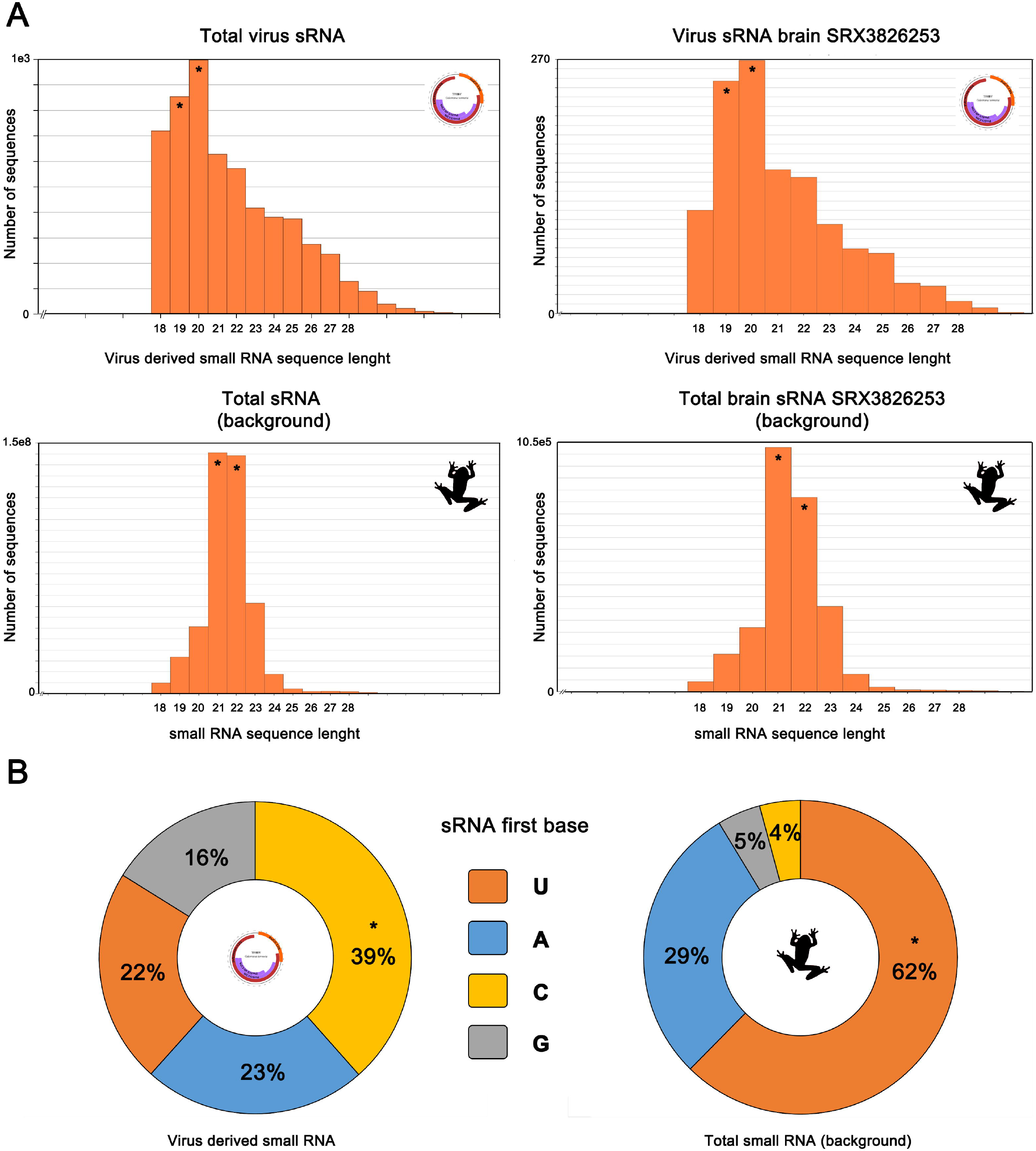
Virus derived small RNA of TFHBV-Ot. (**A**) Total virus derived sRNAs classified by length from bioproject PRJNA437724 (left upper panel) or only from library SRX3826253 (right upper panel) obtained from dissected brain of a female frog. For comparison, the same classification of all sRNA reads is depicted for the complete bioproject (left lower panel) or only from library SRX3826253 (right lower panel). Asterisks denote the two most prevalent sRNA species found in each dataset. (**B**) sRNA first base as percentage of total small RNA reads for the complete bioproject PRJNA437724 (right panel) or only considering virus derived small RNAs (left panel).

**Supplementary Figure 3.**
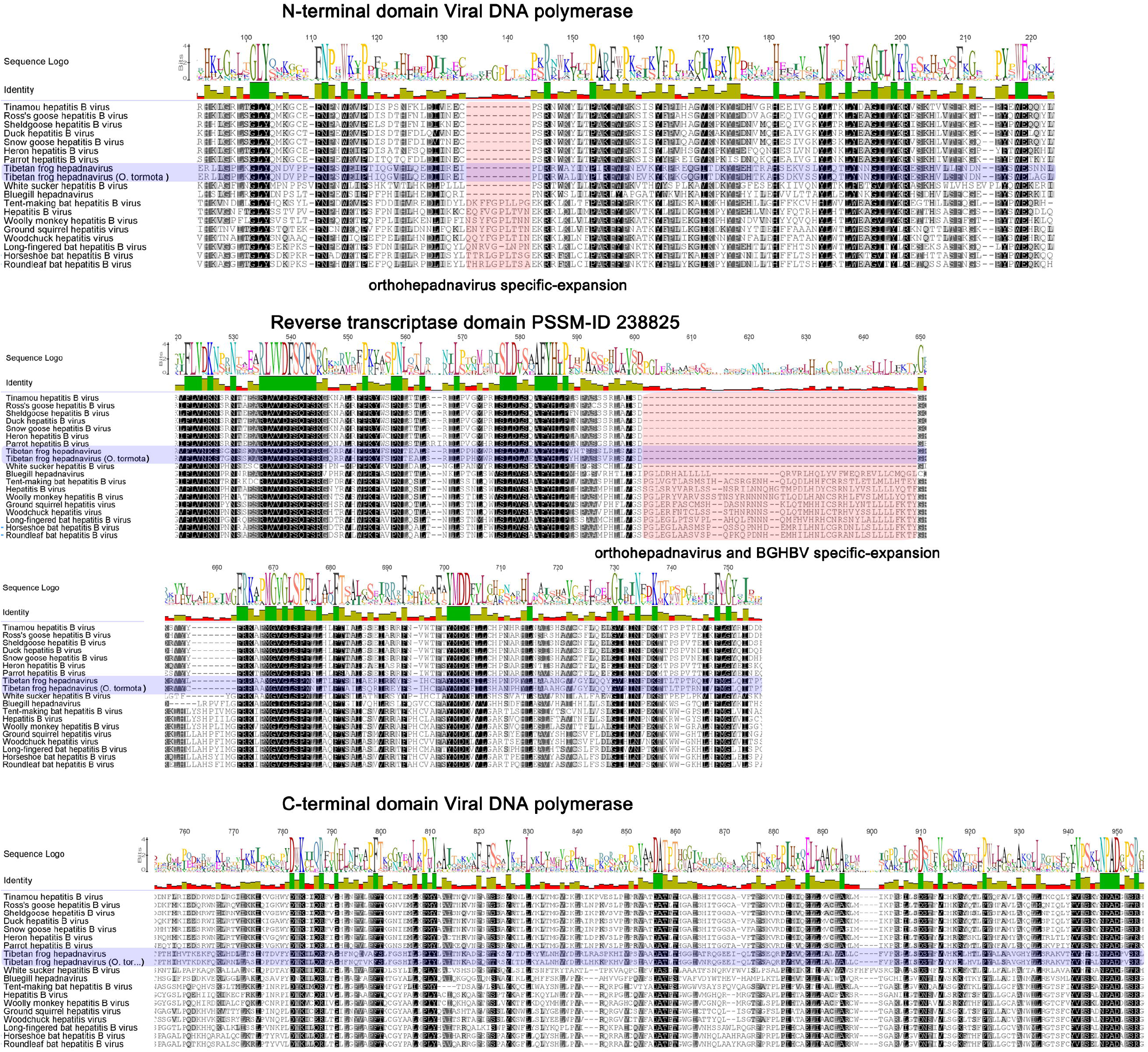
Multiple aa alignment of P proteins of TFHBV-Ot and reported hepadnaviruses generated by MAFTT. Only N/C terminal domains and reverse transcriptase domains are shown. TFHBV are indicated with blue highlighting. Red rectangles depict specific expansions. Similarity is shown from white (low) to black (100% identity).

**Supplementary Table 1.** Tibetan frog hepatitis B virus derived short reads in diverse NGS RNAseq datasets obtained from small RNA samples of dissected tissue/organs from *Odorrana tormota*. Data is single 18-37bp filtered reads obtained with a BGISEQ-500 instrument. Viral reads less than 100 would be treated as typical sequencing bleed over.

| Sample id | Run | BioSample | Experiment | Sex | Tissue | Total reads | Virus reads | Virus RPM |
| --- | --- | --- | --- | --- | --- | --- | --- | --- |
| OT1FBmiRNA | SRR6872795 | SAMN08714513 | SRX3826260 | female | Brain | 28765745 | 650 | 22.6 |
| OT1FMmiRNA | SRR6872796 | SAMN08714514 | SRX3826259 | female | Hindlimb muscle | 28254413 | 948 | 33.6 |
| OT1FSmiRNA | SRR6872797 | SAMN08714515 | SRX3826258 | female | Ovaries | 28222189 | 197 | 7.0 |
| OT1MBmiRNA | SRR6872798 | SAMN08714516 | SRX3826257 | male | Brain | 28399684 | 0 | 0.0 |
| OT1MMmiRNA | SRR6872801 | SAMN08714517 | SRX3826254 | male | Hindlimb muscle | 27133835 | 1 | 0.0 |
| OT1MSmiRNA | SRR6872799 | SAMN08714518 | SRX3826256 | male | Testis | 28887872 | 0 | 0.0 |
| OT2FBmiRNA | SRR6872802 | SAMN08714519 | SRX3826253 | female | Brain | 28005713 | 1233 | 44.0 |
| OT2FMmiRNA | SRR6872804 | SAMN08714520 | SRX3826251 | female | Hindlimb muscle | 26703096 | 2774 | 103.9 |
| OT2FSmiRNA | SRR6872800 | SAMN08714521 | SRX3826255 | female | Ovaries | 26627948 | 1 | 0.0 |
| OT2MBmiRNA | SRR6872803 | SAMN08714522 | SRX3826252 | male | Brain | 26547642 | 0 | 0.0 |
| OT2MMmiRNA | SRR6872791 | SAMN08714523 | SRX3826264 | male | Hindlimb muscle | 28416610 | 0 | 0.0 |
| OT2MSmiRNA | SRR6872792 | SAMN08714524 | SRX3826263 | male | Testis | 29990793 | 0 | 0.0 |
| OT4FMmiRNA | SRR6872793 | SAMN08714525 | SRX3826262 | female | Hindlimb muscle | 29180483 | 0 | 0.0 |
| OT4MMmiRNA | SRR6872794 | SAMN08714526 | SRX3826261 | male | Hindlimb muscle | 28776201 | 0 | 0.0 |

**Supplementary Table 2.** Viruses, GenBank accession numbers, and abbreviations used in this study.

| Virus | GenBank id | Abbreviation |
| --- | --- | --- |
| Bluegill hepatitis B virus | NC_030445.1 | BGHBV |
| Duck hepatitis B virus | MF471768.1 | DHBV |
| Endogenous junco hepatitis B virus | JV160320<sup>a</sup> | eJHBV |
| Ground squirrel hepatitis virus | NC_001484.1 | GSHBV |
| Hepatitis B virus | NC_003977.2 | HBV |
| Heron hepatitis B virus | NC_001486.1 | HHBV |
| Horseshoe bat hepatitis B virus | NC_024444.1 | HBHBV |
| Long-fingered bat hepatitis B virus | NC_020881.1 | LBHBV |
| Parrot hepatitis B virus | NC_016561.1 | PHBV |
| Pomona bat hepatitis B virus | NC_038503.1 | PBHBV |
| Ross's goose hepatitis B virus | NC_005888.1 | RHGBV |
| Roundleaf bat hepatitis B virus | NC_024443.1 | RBHBV |
| Sheldgoose hepatitis B virus | NC_005890.1 | ShHBV |
| Skink hepatitis B virus | SRX213382<sup>b</sup> | SkHBV |
| Snow goose hepatitis B virus | NC_005950.1 | SGHBV |
| Spiny lizard hepatitis B virus | SRX542351<sup>b</sup> | SLHBV-1 |
| Tent-making bat hepatitis B virus | NC_024445.1 | TBHBV |
| Tibetan frog hepatitis B virus (N. parkeri) | NC_030446.1 | TFHBV-Np |
| Tibetan frog hepatitis B virus (O. tormota) | MH700450 | TFHBV-Ot |
| Tinamou hepatitis B virus | NC_035210.1 | THBV |
| White sucker hepatitis B virus | NC_027922.1 | WSHBV |
| Woodchuck hepatitis virus | NC_004107.1 | WHBV |
| Woolly monkey hepatitis B virus | NC_028129.1 | WMHBV |

<sup>a</sup>TSA accession. <sup>b</sup>Accession number of the corresponding SRA library.

## References

1. Dill JA, Camus AC, Leary JH, Di Giallonardo F, Holmes EC, Ng TFF (2016) Distinct viral lineages from fish and amphibians reveal the complex evolutionary history of hepadnaviruses. J Virol JVI: 00832.

2. Lauber, C., Seitz, S., Mattei, S., Suh, A., Beck, J., Herstein, J.,… & Bartenschlager, R (2017) Deciphering the origin and evolution of hepatitis B viruses by means of a family of non-enveloped fish viruses. Cell Host Microbe 22:387–399.

3. Shen JX, Feng AS, Xu ZM, Yu ZL, Arch VS, Yu XJ, Narins PM (2008) Ultrasonic frogs show hyperacute phonotaxis to female courtship calls. Nature 453:914.

4. Shu Y, Xia J, Yu Q, Wang G, Zhang J, He J, Wang H, Zhang L, Wu H (2018) Integrated analysis of mRNA and miRNA expression profiles reveals muscle growth differences between adult female and male Chinese concave-eared frogs (*Odorrana tormota*). Gene doi:10.1016/j.gene.2018.08.007

5. Katoh K, Standley DM (2013) MAFFT multiple sequence alignment software version 7: improvements in performance and usability. Mol Biol Evol 30:772–780.

6. van Hemert FJ, van de Klundert MA, Lukashov VV, Kootstra NA, Berkhout B, Zaaijer HL (2011) Protein X of hepatitis B virus: origin and structure similarity with the central domain of DNA glycosylase. PLoS ONE 6:e23392.

7. Geoghegan JL, Duchêne S, Holmes EC (2017) Comparative analysis estimates the relative frequencies of co-divergence and cross-species transmission within viral families. PLoS Pathog 13:e1006215.

8. Yeh CT, Liaw YF, Ou JH (1990) The arginine-rich domain of hepatitis B virus precore and core proteins contains a signal for nuclear transport. J Virol 64:6141–6147.

9. King AM, Lefkowitz E, Adams MJ, Carstens EB (Eds.) (2011) Virus taxonomy: ninth report of the International Committee on Taxonomy of Viruses. Elsevier.

